# Functional and Genetic Analysis of Viral Receptor ACE2 Orthologs Reveals a Broad Potential Host Range of SARS-CoV-2

**DOI:** 10.1101/2020.04.22.046565

**Authors:** Yinghui Liu, Gaowei Hu, Yuyan Wang, Xiaomin Zhao, Fansen Ji, Wenlin Ren, Mingli Gong, Xiaohui Ju, Yuanfei Zhu, Xia Cai, Jianping Wu, Xun Lan, Youhua Xie, Xinquan Wang, Zhenghong Yuan, Rong Zhang, Qiang Ding

**Author notes:** Corresponding authors (Q.D.); (R.Z.); (Z.Y.). These authors contributed equally to this work (Yinghui Liu, Gaowei Hu, Yuyan Wang, Xiaomin Zhao).

## Abstract

The pandemic of Coronavirus Disease 2019 (COVID-19), caused by severe acute respiratory syndrome coronavirus 2 (SARS-CoV-2), is a major global health threat. Epidemiological studies suggest that bats are the natural zoonotic reservoir for SARS-CoV-2. However, the host range of SARS-CoV-2 and intermediate hosts that facilitate its transmission to humans remain unknown. The interaction of coronavirus with its host receptor is a key genetic determinant of host range and cross-species transmission. SARS-CoV-2 uses angiotensin-converting enzyme 2 (ACE2) as the receptor to enter host cells in a species-dependent manner. It has been shown that human, palm civet, pig and bat ACE2 can support virus entry, while the murine ortholog cannot. In this study, we characterized the ability of ACE2 from diverse species to support viral entry. We found that ACE2 is expressed in a wide range of species, with especially high conservation in mammals. By analyzing amino acid residues of ACE2 critical for virus entry, based on structure of SARS-CoV spike protein interaction with human, bat, palm civet, pig and ferret ACE2, we identified approximately eighty ACE2 proteins from mammals that could potentially mediate SARS-CoV-2 entry. We chose 48 representative ACE2 orthologs among eighty orthologs for functional analysis and it showed that 44 of these mammalian ACE2 orthologs, including those of domestic animals, pets, livestock, and animals commonly found in zoos and aquaria, could bind SARS-CoV-2 spike protein and support viral entry. In contrast, New World monkey ACE2 orthologs could not bind SARS-CoV-2 spike protein and support viral entry. We further identified the genetic determinant of New World monkey ACE2 that restricts viral entry using genetic and functional analyses. In summary, our study demonstrates that ACE2 from a remarkably broad range of species can facilitate SARS-CoV-2 entry. These findings highlight a potentially broad host tropism of SARS-CoV-2 and suggest that SARS-CoV-2 might be distributed much more widely than previously recognized, underscoring the necessity to monitor susceptible hosts to prevent future outbreaks.

## Introduction

Coronaviruses are a group of positive-stranded, enveloped RNA viruses that circulate broadly among humans, other mammals, and birds, causing respiratory, enteric, or hepatic diseases^1^. In the last two decades, coronaviruses have caused three major outbreaks: severe acute respiratory syndrome (SARS), Middle East respiratory syndrome (MERS) and the recent Coronavirus Disease 2019 (COVID-19)^2,3^. As of April 24, 2020, COVID-19 has already caused 2.7 million infections, leading to 192,000 deaths globally. The pathogen responsible is a novel coronavirus-severe acute respiratory syndrome coronavirus 2 (SARS-CoV-2) ^4,5^. Phylogenetic and epidemiological analyses suggest that SARS-CoV, MERS-CoV and SARS-CoV-2 likely originated from bats, with SARS-CoV spreading from bats to palm civets to humans, and MERS-CoV spreading from bats to camel to humans^6^. However, the intermediate host of SARS-CoV-2, fueling spillover to humans, remains unknown.

The SARS-CoV-2 genome encodes a spike (S) protein, the receptor-binding domain (RBD) of which binds the cellular receptor angiotensin-converting enzyme 2 (ACE2) to mediate viral entry^5,7^. Following binding of ACE2, the S protein is subsequently cleaved by the host transmembrane serine protease 2 (TMPRSS2) to release the spike fusion peptide, promoting virus entry into target cells^7,8^. It has been repeatedly demonstrated that the interaction of a virus with (a) species-specific receptor(s) is a primary determinant of host tropism and therefore constitutes a major interspecies barrier at the level of viral entry^9^. For example, murine ACE2 does not efficiently bind the SARS-CoV or SARS-CoV-2 S protein, hindering viral entry into murine cells; consequently, a human ACE2 transgenic mouse was developed as an *in vivo* model to study the infection and pathogenesis of these two viruses^10,11^.

ACE2 is expressed in a diverse range of species throughout the subphylum *Vertebrata*. Several recent studies demonstrated that ferrets, cats, dogs and some non-human primates are susceptible to SARS-CoV-2^12–16^. However, the exact host tropism of SARS-CoV-2 remains unknown and it is urgent to identify the putative zoonotic reservoirs to prevent future outbreak. To address this question, we systematically analyzed ACE2 orthologs from a broad range of species for their ability to support SARS-CoV-2 entry. Our data demonstrate that an evolutionarily diverse set of ACE2 species variants can mediate SARS-CoV-2 glycoprotein-dependent entry, suggesting that SARS-CoV-2 has a broad host range at the level of virus entry that may contribute to cross-species transmission and viral evolution.

## RESULTS

### Evolutionary and phylogenetic analyses of ACE2 orthologs from a diversity of species

ACE2, the cellular receptor for SARS-CoV-2 and SARS-CoV, is expressed in a diverse range of vertebrate animals. We analyzed the protein sequences of 294 ACE2 orthologs in the NCBI database, from birds (75 species), alligators (4 species), turtles (4 species), lizards (9 species), mammals (129 species), amphibians (4 species), coelacanths (1 species), bone fish (67 species) and cartilaginous fish (1 species) (Fig. S1). These ACE2 orthologs range from 344 to 861 amino acid residues in length and are characterized by an N-terminal leucine-rich repeat (LRR) domain and a C-terminal low-complexity acidic region (LCAR). Structures of SARS-CoV S protein complexed with human ACE2 or other susceptible orthologs have been solved, and five critical, highly conserved amino acid residues of ACE2 have been identified that are indispensable for interaction with S protein and viral entry^8,17–19^. Based on this structural information and conservation of these 5 critical residues, we carried out primary sequence alignment across the 294 ACE2 proteins (Fig. S1). Our analysis revealed that ACE2 orthologs from 80 species that could potentially function as SARS-CoV-2 receptors (Fig. S1, and Fig. 1). All of the 80 ACE2 orthologs were derived from mammals, including wild animals, domestic animals, pets, livestock and even animals in the zoo or aquaria, with protein size ranging from 790 to 811 amino acids (Table S1). Other ACE2 orthologs from mammals (49/129 species) that were predicted as non-functional receptors of SARS-CoV and SARS-CoV-2 are summarized in Table S2 and were not included in the experiments described in the present study. Interestingly, species in Table S2, such as Chinese tree shrew and guinea pigs were not susceptible to SARS-CoV-2 infection as demonstrated by other studies ^20,21^.

**Figure 1.**
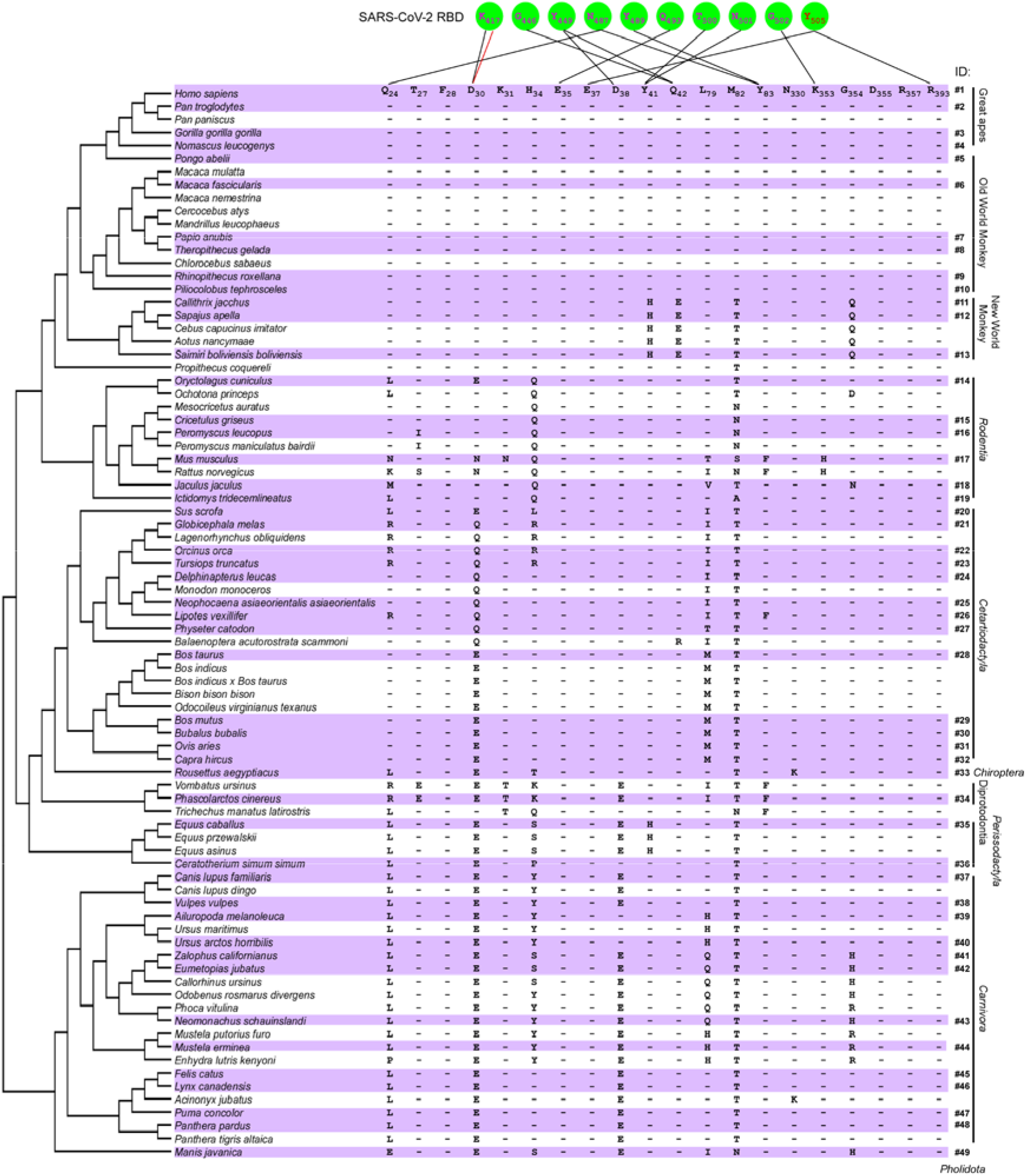
Phylogenetic analysis of ACE2 orthologs with potential to support SARS-CoV-2 entry and alignment of ACE2 residues at the interface with the viral spike protein. The ACE2 protein sequences (Supplemental Table 1), as well as *Mus musculus* (mouse) and *Rattus norvegicus* (rat) ACE2, were chosen and analyzed by MEGA-X (Version 10.05) software and MUSCLE algorithm. The phylogenetic tree was built using Neighbor Joining method of the MEGA-X. The contacting residues of human ACE2 (distance cutoff of 4 Å) at the SARS-CoV-2 receptor binding domain (RBD)/ACE2 interface are shown. The contacting network involves at least 20 residues in ACE2 and 10 residues in the SARS-CoV-2 RBD, which are listed and connected by a solid line. Black lines indicate hydrogen bonds, and the red line represents a salt-bridge interaction. The tested species are highlighted in purple and the ID number of each species in subsequent experiments is labeled on the right. Only the amino acids different from human are shown.

We performed phylogenetic analysis of these 80 ACE2 orthologs to explore their potential function in mediating virus infection and to gain insights into the evolution of the ACE2 protein (Fig.1, left panel). Additionally, we aligned the twenty residues of ACE2 located at the interface with the SARS-CoV-2 S protein^22–24^ (Fig. 1, right panel). The ACE2 protein sequences were highly conserved across the species we analyzed. Of note, the twenty residues at the ACE2-S protein interface were identical across the Catarrhini, which includes great apes and Old World monkeys. However, these residues in the ACE2 orthologs of New World monkeys were less conserved. For example, Y41 and Q42 in human ACE2 are responsible for the formation of hydrogen bonds with S protein and are highly conserved across all other species but are substituted by H and E, respectively in New World monkeys. In non-primate mammals, an increasing number of substitutions are evident, even in residues such as Q24, D30, D38, and Y83 that form hydrogen bonds or salt-bridges with S protein (Fig. 1).

Collectively, our analysis suggests that ACE2 orthologs are highly conserved across a wide range of mammals, and many of these ACE2 orthologs might function as an entry receptor for SARS-CoV-2.

### Interaction of ACE2 proteins with SARS-CoV-2 spike protein

Based on our evolutionary analysis, we chose 48 representative ACE2 orthologs (Table S1 and Fig. 1) from *Primates, Rodentia, Cetartiodactyla, Chiroptera, Diprotodontia, Perissodactyla, Carnivora* and *Pholidota* for further analysis. We assessed whether they support SARS-CoV-2 entry by ectopically expressing each ortholog in HeLa cells, which have limited endogenous ACE2 expression^5^. These 48 species include wild animals, animals in the zoo and aquaria, pets and livestock frequently in close contact with humans, model animals used in biomedical research, and endangered species (Fig. 1).

Binding of SARS-CoV-2 S protein to ACE2 is a prerequisite for viral entry. To examine this, we employed a cell-based assay that used flow cytometry to assess the binding of S protein to different ACE2 orthologs. We cloned the cDNA of 49 ACE2 orthologs (murine ACE2 was included as a negative control), each with a C-terminal FLAG tag, into a bicistronic lentiviral vector (pLVX-IRES-zsGreen1) that expresses the fluorescent protein zsGreen1 via an IRES element and can be used to monitor transduction efficiency. Next, a purified fusion protein consisting of the S1 domain of SARS-CoV-2 S protein and an Fc domain of human lgG (S1-Fc) was incubated with HeLa cells transduced with the ACE2 orthologs. Binding of S1-Fc to ACE2 was then quantified by flow cytometry as the percent of cells positive for S1-Fc among the ACE2 expressing cells (zsGreen1^+^ cells). As expected, the binding of S1-Fc to HeLa cells expressing mouse ACE2 was very low and comparable to that of the empty vector control, whereas S1-Fc protein efficiently bound to HeLa cells expressing human ACE2, which is consistent with previous reports^5,8^. Surprisingly, we found that ACE2 from 44/49 species could bind the S1-Fc protein, albeit slightly less efficiently than human ACE2 (Fig. 2A). In contrast, ACE2 from *Callithrix jacchus* (marmoset, #11), *Sapajus apella* (tufted capuchin, #12), and *Saimiri boliviensis boliviensis* (squirrel monkey, #13)—all New World monkeys—failed to bind S1-Fc; ACE2 from *Phascolarctos cinereus* (koala, #34) and *Mustela ermine* (stoat, #44) bound only poorly to the S1-Fc fusion (Fig. 2).

**Figure 2.**
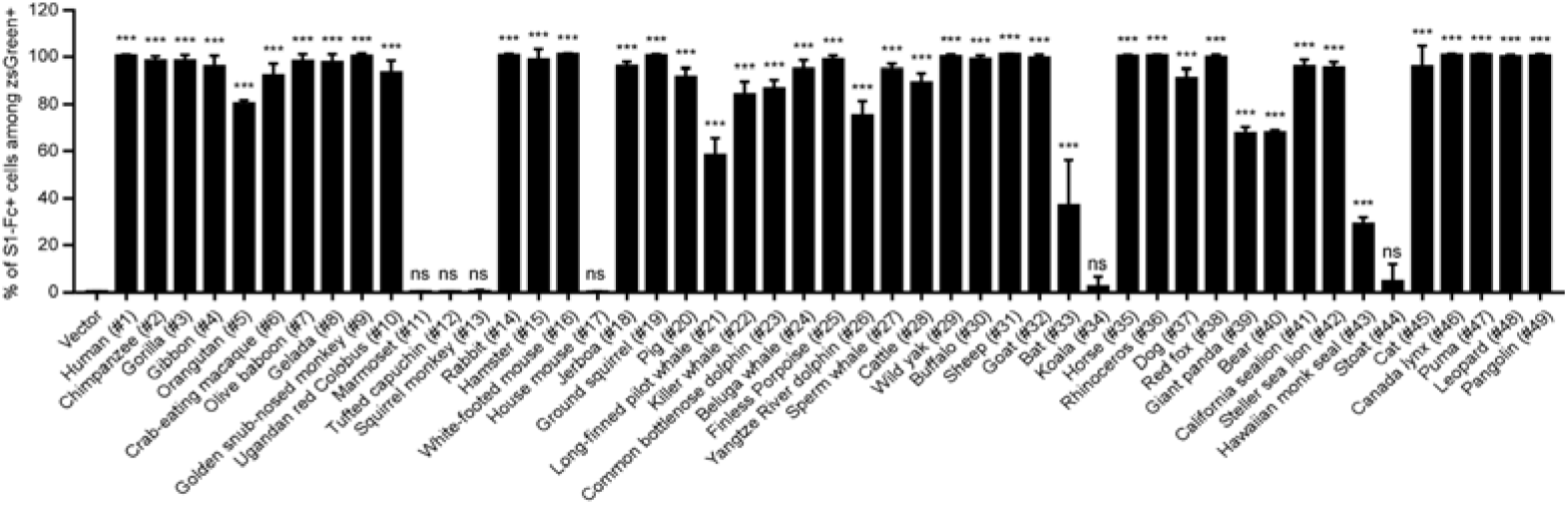
Binding of the SARS-CoV-2 spike protein to different ACE2 orthologs. HeLa cells were transduced with ACE2 orthologs of the indicated species, incubated with the recombinant S1 domain of SARS-CoV-2 spike protein C-terminally fused with Fc, and then stained with goat anti-human IgG (H + L) conjugated to Alexa Fluor 647 for flow cytometry analysis. Values are expressed as the percent of cells positive for S1-Fc among the ACE2 expressing cells (zsGreen1+cells) and are means plus standard deviations (SD) (error bars). ns, no significance; ***, P < 0.001. Significance assessed by one-way ANOVA.

In summary, these results demonstrate that ACE2 proteins from a broad range of diverse species can bind to the SARS-CoV-2 S protein, suggesting that these species may indeed be capable of mediating viral uptake.

### Functional assessment of ACE2 orthologs in SARS-CoV-2 entry

It has been shown that HeLa cells lacking expression of endogenous ACE2 were not permissive to SARS-CoV-2 infection^5^. To test directly whether different ACE2 orthologs can indeed mediate viral entry, we performed genetic complementation experiments in HeLa cells.

HeLa cells ectopically expressing individual ACE2 orthologs were infected with SARS-CoV-2 (MOI=1). At 48 h post-infection, the complemented HeLa cells were fixed and underwent immunofluorescent staining for intracellular viral nucleocapsid protein, an indicator of virus replication. As expected, HeLa cells expressing mouse ACE2 were not permissive to SARS-CoV-2 infection while those expressing human ACE2 were permissive (Fig.3, #17 and #1, respectively). Consistent with our binding data, HeLa cells expressing ACE2 orthologs from marmoset (#11), tufted capuchin (#12), squirrel monkey (#13) or koala (#34) were non-permissive to SARS-CoV-2 infection. HeLa cells expressing ACE2 from stoat (#44) were permissive, albeit with low efficiency; the remaining 44 ACE2 orthologs supported SARS-CoV-2 infection, as evidenced by readily detectable viral nucleocapsid protein within ACE2-expressing (zsGreen1+) cells (Fig. 3).

**Figure 3.**
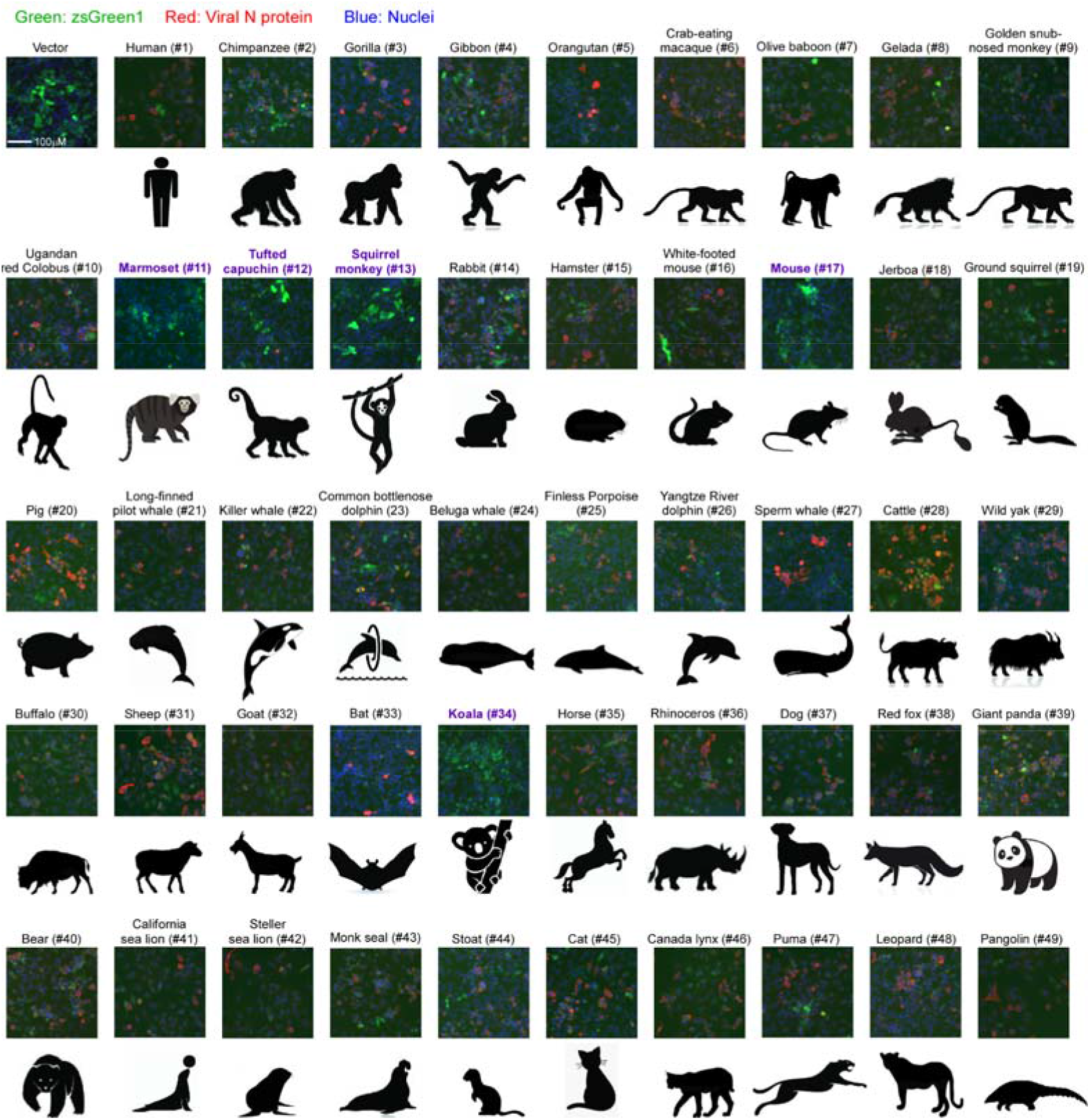
Functional assessment of ACE2 orthologs mediating SARS-CoV-2 virus entry. HeLa cells transduced with lentiviruses expressing ACE2 orthologs or empty vector were infected with SARS-CoV-2 virus (MOI=1). Expression of the viral nucleocapsid protein was visualized by immunofluorescence microscopy. Viral nucleocapsid (N) protein (red) and nuclei (blue) are shown. Green signal indicates the transduction efficiency of ACE2 orthologs. Marmoset (#11), tufted capuchin (#12), squirrel monkey (#13), and koala (#34) were non-permissive to SARS-CoV-2 infection, highlighted in purple. The images were merged and edited using Image J software.

Collectively, our results demonstrate that SARS-CoV-2 can utilize ACE2 from evolutionarily diverse species of mammals as a cellular receptor for viral entry, suggesting that SARS-CoV-2 may have a broad host range.

### The genetic determinants of ACE2 from New World monkeys that restrict SARS-CoV-2 entry

Although the overall protein sequences of ACE2 were largely conserved across all tested species (71%–100% identity compared with human ACE2) (Fig.S2), our data showed that high sequence identity does not necessarily correlate with its function to support virus entry. For example, as shown in Fig. 3 and Fig. 4, ACE2 orthologs from the New World monkeys marmoset (#11), tufted capuchin (#12), and squirrel monkey (#13) had limited or undetectable ability to mediate SARS-CoV-2 entry despite sharing 92-93% identity with human ACE2. In contrast, the ACE2 proteins from *Bos taurus* (cattle, #28) or *Sus scrofa* (pig, #20) efficiently facilitated virus entry, even with 78% or 81% identity, respectively, to human ACE2 (Fig. S2). Thus, we hypothesized that changes in critical residues in ACE2 proteins from New World monkeys may restrict viral entry.

**Figure 4.**
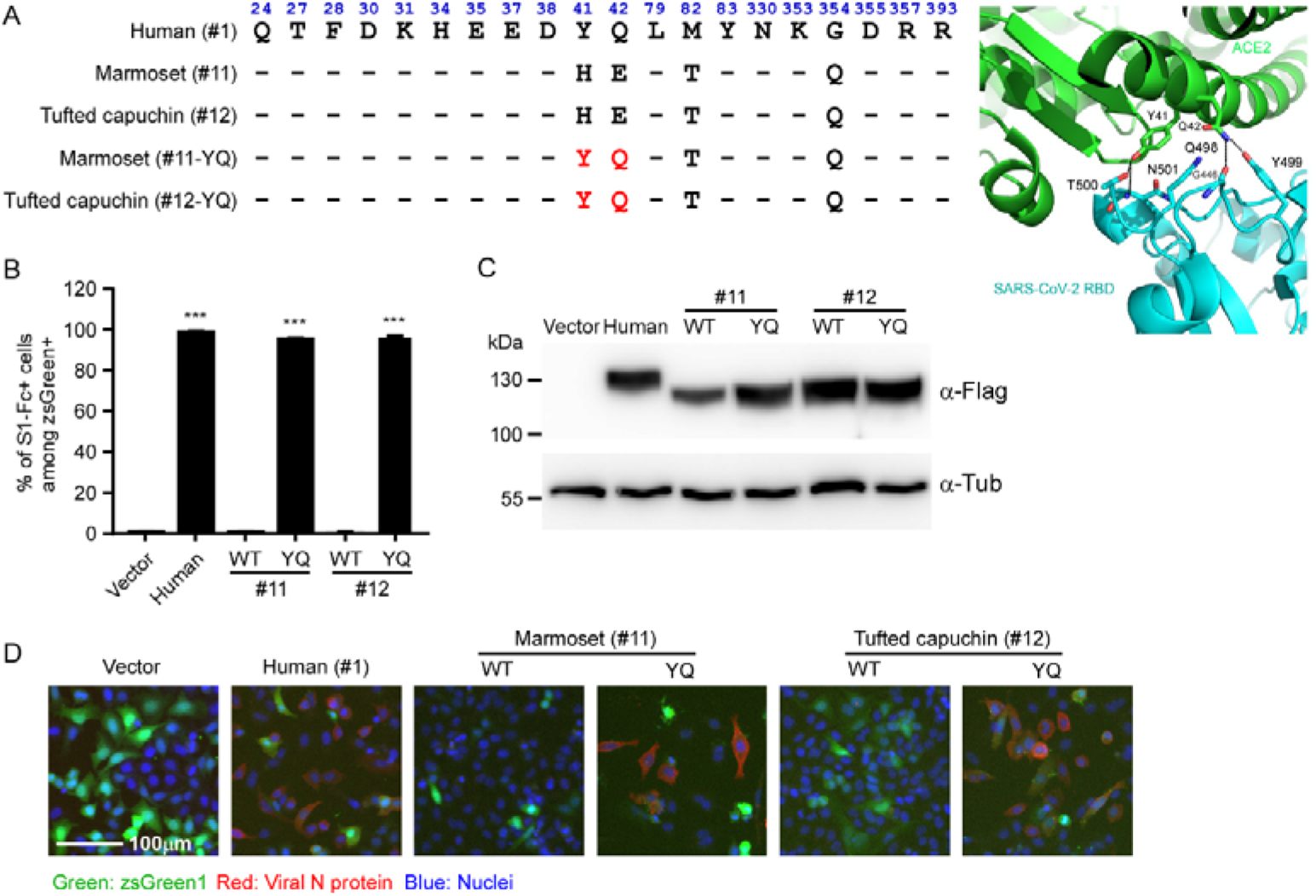
Identification of the species-specific genetic determinants of ACE2 restriction of SARS-CoV-2 entry. (A) Left panel: Alignment of the contacting residues of human ACE2 (distance cutoff of 4 Å) at the SARS-CoV-2 receptor binding domain (RBD)/ACE2 interface with orthologs from the New World monkeys marmoset (#11) and tufted capuchin (#12). The mutations introduced into marmoset (#11) and tufted capuchin (#12) ACE2 at position 41 and 42 are indicated in red. Right panel: The binding interface of human ACE2 with SARS-CoV-2 receptor-binding domain (RBD) surrounding ACE2 Y41 and Q42. Residue Y41 forms hydrogen bonds with T500 and N501 of SARS-CoV-2 RBD, and Q42 can also interact with G446 or Y449 by hydrogen bonds. The differences in ACE2 from New World monkeys, especially the Y41H replacement, may disrupt the hydrogen-bonding interactions and impair the binding with SARS-CoV-2 spike. PDB code of the complex of human ACE2 with SARS-CoV-2: 6M0J. (B-C) HeLa cells transduced with ACE2 orthologs of the indicated species or mutants were incubated with the recombinant S1 domain of SARS-CoV-2 spike C-terminally fused with Fc to determine the binding of ACE2 with SARS-CoV-2 spike as described in Fig. 2A and B. Values are means plus standard deviations (SD) (error bars). ns, no significance; ***, P < 0.001. Significance assessed by one-way ANOVA. (D) HeLa cells transduced with lentiviruses expressing ACE2 orthologs (or mutants) or empty vector were infected with SARS-CoV-2 virus (MOI=1). The infection was determined by immunofluorescence microscopy as described in Fig.3. The images were merged and edited using Image J software.

New World monkeys are widely used in biomedical research. Our results showed that their ACE2 proteins do not bind SARS-CoV-2 S protein and do not promote virus entry, which is in line with a recent finding that marmosets are resistant to SARS-CoV-2 infection^13^. To identify the genetic determinants within ACE2 orthologs from New World monkeys that restrict viral entry, we first analyzed the ACE2 protein residues that contact the S protein, especially those that form hydrogen bonds or salt bridges with S protein, such as Q24, D30, E35, E37, D38, Y41, Q42, Y83, K353 and R393^22–24^. When comparing with orthologs that support SARS-CoV-2 entry, we found that residues at the ACE2-S interface in New World monkeys only differed at H41 and E42 (Fig.1 and Fig. 4A). The hydroxyl group of the Tyr (Y) at human ACE2 position 41 forms hydrogen bonds with the side chain oxygen atom of T500 and side chain nitrogen atom of N501 in the SARS-CoV-2 S protein. The side chain nitrogen atom of Q42 of human ACE2 forms hydrogen bonds with the main chain oxygen atom of G446 and side chain hydroxyl group of Y449 of the SARS-CoV-2 S protein. Changes at these two consecutive residues, 41 and 42, may disrupt critical hydrogen-bonding interactions and thus impair the binding of New World monkey ACE2 with SARS-CoV-2 S protein (Fig. 4A, right panel).

To directly uncover the molecular basis for the inability of New World monkey ACE2 to function as a SARS-CoV-2 receptor, we humanized marmoset and tufted capuchin ACE2 by mutating the ACE2 orthologs at position 41 and 42 into Y and Q, respectively (Fig.4A). These humanized orthologs were then transduced into HeLa cells to assess binding with the SARS-CoV-2 spike protein. Remarkably, both the humanized marmoset and tufted capuchin ACE2 orthologs were now able to bind the SARS-CoV-2 S protein with efficiency comparable to that of human ACE2 (Fig. 4B and C). To confirm whether this gain-of-function phenotype was functional in the context of infection, HeLa cells ectopically expressing WT or these humanized ACE2 orthologs were infected with SARS-CoV-2 (MOI=1). At 48 h post-infection, the complemented HeLa cells were subjected to immunofluorescent staining for intracellular viral nucleocapsid protein. As we observed before and consistent with our binding data, HeLa cells expressing ACE2 orthologs from marmoset (#11) or tufted capuchin (#12) were non-permissive to SARS-CoV-2 infection (Fig. 4D). However, the humanized ACE2 orthologs from marmoset (#11-YQ) or tufted capuchin (#12-YQ) rendered the HeLa cells permissiveness to infection (Fig. 4D), demonstrating that altering the residues at position 41 and 42 into human counterparts confers the ability of ACE2 orthologs from New World monkeys of binding to SARS-CoV-2 spike protein and mediating viral entry, thereby determining the ability of these ACE2 proteins to be used as viral receptors.

Thus, our analysis identifies the genetic determinants of ACE2 in New World monkeys necessary for the protein’s function as the SARS-CoV-2 cellular receptor and provides greater insight into the species-specific restriction of viral entry, which can inform the development of animal models.

## Discussion

To prevent the zoonotic transmission of SARS-CoV-2 to humans, the identification of animal reservoirs or intermediate hosts of SARS-CoV-2 is of great importance. Recently, a coronavirus was identified in pangolins with 90% sequence identity compared to SARS-CoV-2^25,26^. However, the result of such phylogenetic analysis does not necessarily support the notion that pangolins are indeed an intermediate host of SARS-CoV-2. The host range and animal reservoirs of SARS-CoV-2 remain to be explored.

For the cross-species transmission of SARS-CoV-2 from intermediate hosts to humans, the virus needs to overcome at least two main host genetic barriers: the specificity of the viral S protein-ACE2 receptor interactions and the ability to escape the host’s antiviral immune response. The interaction of a virus with its host cell receptor is the first step to initiate virus infection and is a critical determinant of host species range and tissue tropism. SARS-CoV-2 uses the cellular receptor ACE2 in a species-specific manner: human, palm civet, bat and pig ACE2 can support virus entry whereas mouse ACE2 cannot^5^. To explore possible SARS-CoV-2 animal reservoirs and intermediate hosts, we analyzed ACE2 genes from 294 vertebrates, particularly mammals. Our results suggest that ACE2 orthologs are largely conserved across vertebrate species, indicating the importance of its physiological function. Notably, we also found that ACE2 orthologs from a wide range of mammals could act as a functional receptor to mediate SARS-CoV-2 infection when ectopically expressed in HeLa cells, suggesting that SARS-CoV-2 may have a diverse range of hosts and intermediate hosts.

It is of note that our findings are based on a functional study of ACE2 proteins during authentic SARS-CoV-2 infection instead of using pseudotyped virus. Our results are consistent with recent *in vivo* findings that ferrets, cats, dogs, and Old World monkeys are susceptible to SARS-CoV-2 infection but not marmosets, which are New World monkeys^13–15^. The host range or specificity of a virus is often limited due to several reasons, such as the lack of host factors the virus depends on or the incompatibility of these factors’ orthologs in different species. Alternatively, but not necessarily mutually exclusive, the ability to evade the antiviral immune response of a given host can also shape the species tropism of viruses^9,27^.

Development of prophylactic vaccines or effective antivirals are urgently needed to combat SARS-CoV-2 infection^28^. Establishment of better animal models to evaluate the efficacy of vaccine candidates and antiviral strategies *in vivo* is thus of utmost importance. Additionally, there is a need for suitable, experimentally tractable animal models to dissect mechanistically viral transmission and pathogenesis. Human ACE2 transgenic mice have been used to study SARS-CoV and SARS-CoV-2 *in vivo*^10,11^. However, the unphysiologically high expression level of ACE2 driven by the ubiquitous K14 promoter may not recapitulate the human disease caused by SARS-CoV-2. Recently, a ferret model of SARS-CoV-2 infection was established that mimics transmission and recapitulates aspects of human disease^12^. Our study found that ACE2 from multiple species of laboratory animals, including but not limited to ferrets, crab-eating macaques, and Chinese hamsters, could be utilized by SARS-CoV-2 to mediate viral infection. Our data provide a rationale to assess the susceptibility of such species whose ACE2 ortholog serves as a functional receptor for SARS-CoV-2. Our results further demonstrate that ACE2 orthologs from three New World monkey species (marmoset (#11), tufted capuchin (#12), and squirrel monkey (#13)) do not support SARS-CoV-2 entry. We identified specific residues—H41 and E42—within ACE2 that restrict SARS-CoV-2 in these species. Substituting these critical amino acids with those found in human ACE2 rendered these ACE2 orthologs able to support SARS-CoV-2 entry.

Our unexpected finding that SARS-CoV-2 uses ACE2 from diverse species highlights the importance of surveilling animals in close contact with humans as potential zoonotic reservoirs. We found that pets such as cats and dogs, livestock such as pigs, cattle, rabbits, sheep, horses, and goats, and even some animals kept frequently in zoos or aquaria may serve as intermediate hosts for virus transmission. Our study also identified a broad range of wild animals as potential susceptible hosts of SARS-CoV-2, highlighting the importance of banning illegal wildlife trade and consumption.

In summary, ours is the first study to systematically assess the functionality of ACE2 orthologs from nearly 50 mammalian hosts using the authentic SARS-CoV-2 virus, which provides new insight into the potential host range and cross-species transmission of this virus. It also suggests that SARS-CoV-2 might be much more widely distributed than previously thought, underscoring the necessity of monitoring susceptible hosts, especially their potential for causing zoonosis to prevent future outbreaks.

## ACKNOWLEDGEMENTS

We thank Drs. Alexander Ploss (Princeton University), Jin Zhong (Institut Pasteur of Shanghai, CAS), Ke Lan (Wuhan University), Chunliang Xu (Albert Einstein College of Medicine) and Jenna M. Gaska for suggestions and revision of the manuscript. We wish to acknowledge Di Qu, Zhiping Sun, Wendong Han and other colleagues at the Biosafety Level 3 Laboratory of Fudan University for help with experiment design and technical assistance. We are grateful to Yingjie Zhang and Ruiqi Chen (Tsinghua University) for validating gene sequences.

This work was supported by Tsinghua-Peking University Center of Life Sciences (045-61020100120), National Natural Science Foundation of China (32041005), Tsinghua University Initiative Scientific Research Program (2019Z06QCX10), Beijing Advanced Innovation Center for Structure Biology (100300001), Start-up Foundation of Tsinghua University (53332101319), Shanghai Municipal Science and Technology Major Project (20431900400) and Project of Novel Coronavirus Research of Fudan University.

## MATERIALS AND METHODS

### Cell cultures and SARS-CoV-2 virus

HEK293T cells (American Tissue Culture Collection, ATCC, Manassas, VA, CRL-3216), Vero E6 (Cell Bank of the Chinese Academy of Sciences, Shanghai, China) and HeLa (ATCC #CCL-2) were maintained in Dulbecco’s modified Eagle medium (DMEM) (Gibco, NY, USA) supplemented with 10% (vol/vol) fetal bovine serum (FBS), 10mM HEPES, 1mM sodium pyruvate, 1×non-essential amino acids, and 50 IU/ml penicillin/streptomycin in a humidified 5% (vol/vol) CO2 incubator at 37°C. Cells were tested routinely and found to be free of mycoplasma contamination. The SARS-CoV-2 strain nCoV-SH01 (GenBank accession no. MT121215) was isolated from a COVID-19 patient and propagated in Vero E6 cells for use. All experiments involving virus infections were performed in the biosafety level 3 facility of Fudan University following the regulations.

### Plasmids

The cDNAs encoding ACE2 orthologs (Table S1) were synthesized by GenScript and cloned into pLVX-IRES-zsGreen1 vectors (Catalog No. 632187, Clontech Laboratories, Inc) with a C-terminal FLAG tag. ACE2 mutants were generated by Quikchange (Stratagene) site-directed mutagenesis. All of the constructs were verified by Sanger sequencing.

### Lentivirus production

Vesicular stomatitis virus G protein (VSV-G) pseudotyped lentiviruses expressing ACE2 orthologs tagged with FLAG at the C-terminus were produced by transient co-transfection of the third-generation packaging plasmids pMD2G (Addgene #12259) and psPAX2 (Addgene #12260) and the transfer vector with VigoFect DNA transfection reagent (Vigorous) into HEK293T cells. The medium was changed 12 h post transfection. Supernatants were collected at 24 and 48h after transfection, pooled, passed through a 0.45-μm filter, and frozen at −80°C.

### Phylogenetic analysis and sequence alignment

The amino acid sequences of ACE2 orthologs for jawed vertebrates (Gnathostomata) were exported from the NCBI nucleotide database. Numbers in each sequence correspond to the GenBank accession number. 81 sequences were collected for the presence of five critical viral spike-contacting residues of ACE2 corresponding to amino acids Lys31, Glu35, Asp38, Met82 and Lys353 in human ACE2 (NM_001371415.1). The protein sequences of different species were then passed into MEGA-X (Version 10.05) software for further analysis. The alignment was conducted using the MUSCLE algorithm ^29^. Then the alignment file was used to construct the phylogenetic tree (Neighbor Joining option of the MEGA-X with default parameter).

### Western blotting

Sodium dodecyl sulfate-polyacrylamide gel electrophoresis (SDS-PAGE) immunoblotting was performed as follows: After trypsinization and cell pelleting at 2,000 × g for 10 min, whole-cell lysates were harvested in RIPA lysis buffer (50 mM Tris-HCl [pH 8.0], 150mM NaCl, 1% NP-40, 0.5% sodium deoxycholate, and 0.1% SDS) supplemented with protease inhibitor cocktail (Sigma). Lysates were electrophoresed in 12% polyacrylamide gels and transferred onto nitrocellulose membrane. The blots were blocked at room temperature for 0.5 h using 5% nonfat milk in 1× phosphate-buffered saline (PBS) containing 0.1% (v/v) Tween 20. The blots were exposed to primary antibodies anti-β-Tubulin (CW0098, CWBIO), or anti-FLAG (F7425, Sigma) in 5% nonfat milk in 1× PBS containing 0.1% Tween 20 for 2 h. The blots were then washed in 1× PBS containing 0.1% Tween 20. After 1h exposure to HRP-conjugated secondary antibodies, subsequent washes were performed and membranes were visualized using the Luminescent image analyzer (GE).

### Surface ACE2 binding assay

HeLa cells were transduced with lentiviruses expressing the ACE2 from different species for 48 h. The cells were collected with TrypLE (Thermo #12605010) and washed twice with cold PBS. Live cells were incubated with the recombinant protein, S1 domain of SARS-CoV-2 spike C-terminally fused with Fc (Sino Biological #40591-V02H, 1μg/ml) at 4 °C for 30 min. After washing, cells were stained with goat anti-human IgG (H + L) conjugated with Alexa Fluor 647 (Thermo #A21445, 2 μg/ml) for 30 min at 4 °C. Cells were then washed twice and subjected to flow cytometry analysis (Thermo, Attune™ NxT).

### Immunofluorescence staining of viral nucleocapsids

HeLa cells were transduced with lentiviruses expressing the ACE2 from different species for 48 h. Cells were then infected with nCoV-SH01 at an MOI of 1 for 1 h, washed three times with PBS, and incubated in 2% FBS culture medium for 48 h for viral antigen staining. Cells were fixed with 4% paraformaldehyde in PBS, permeablized with 0.2% Triton X-100, and incubated with the rabbit polyclonal antibody against SARS-CoV nucleocapsid protein (Rockland, 200-401-A50, 1μg/ml) at 4 °C overnight. After three washes, cells were incubated with the secondary goat anti-rabbit antibody conjugated with Alexa Fluor 488 (Thermo #A11034, 2 μg/ml) for 2 h at room temperature, followed by staining with 4’,6-diamidino-2-phenylindole (DAPI). Images were collected using an EVOS™ Microscope M5000 Imaging System (Thermo #AMF5000). Images were processed using the ImageJ program (http://rsb.info.nih.gov/ij/).

### Statistics analysis

One-way analysis of variance (ANOVA) with Tukey’s honestly significant difference (HSD) test was used to test for statistical significance of the differences between the different group parameters. *P* values of less than 0.05 were considered statistically significant.

#### SUPPLEMENTAL INFORMATION

**Supplemental Figure 1.**
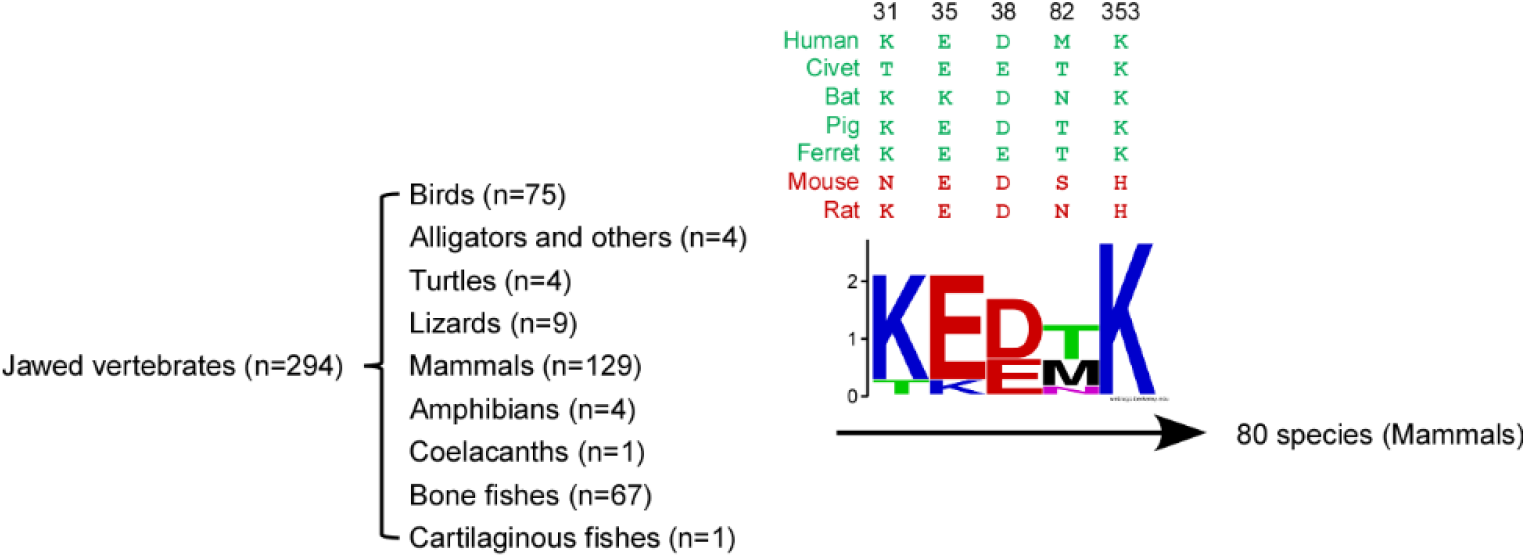
ACE2 orthologs from the jawed vertebrates. ACE2 orthologs were recorded in the NCBI dataset and further parsed to 80 ACE2 orthologs with potential function for supporting SARS-CoV-2 entry based on conservation of the 5 amino acids required for binding between the host receptor ACE2 and the SARS-CoV spike protein ^8,17–19^.

**Supplemental Figure 2.**
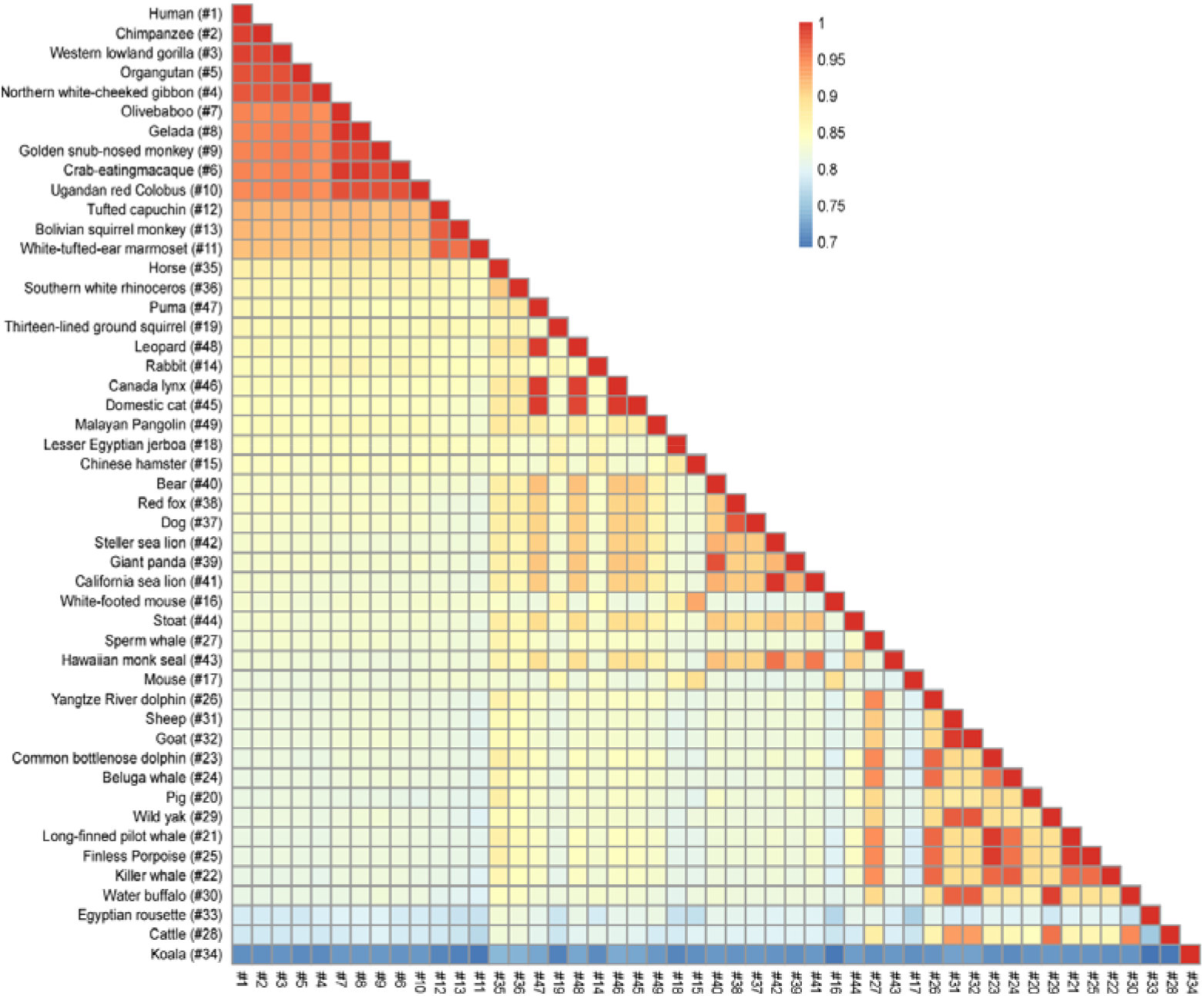
Protein sequence identity matrices of ACE2 from the tested species. The ACE2 sequences from different species were analyzed using SIAS (Sequence Identity And Similarity) tool (http://imed.med.ucm.es/Tools/sias.html) to determine the percent identity of ACE2 proteins across different species.

